# Perception and Readiness Towards Inter-Professional Education Among Different Health Care Disciplines at Khyber Medical University Peshawar

**DOI:** 10.1101/2022.12.21.521507

**Authors:** Yaser Ud-din, Nasreen Ghani, Zubaida Khatoon, Zartasha, Haidar Ali, Sadar Badshah, Syed Hassan, Mehboob Ali, Naveed Iqbal, Abdul Hameed, Qaisar Shehzad

## Abstract

Interprofessional learning (IPL) is an approach that teaches students diverse disciplines to communicate with each other about their professional knowledge in order to acquire a more complex one understanding of the current situation. According to WHO definition of IPE “it’s a process between two or more professionals learn about from, and with one another to permit effective association and enhance health outcome. The aim of this study was to measure perception and readiness towards IPE among different health care disciplines. A cross-sectional study was done with two hundred and eighteen post graduate Nursing, public health, physiotherapy, and basic medical sciences students from September,2020 to January,2021 at Khyber Medical University Peshawar Pakistan. Convenient sampling technique was used to collect data. The Readiness for Inter-Professional Learning Scale (RIPLS) and Interdisciplinary Education Perception Scale (IEPS) were used to measure the readiness and perception of students regarding inter-professional leaning. The data were analyzed using software Statistical Package for Social Science (SPSS) (Version 23). The RIPLS was completed by a total of 218 participants (response rate 100%, 61 Nursing students 28%, 51 physiotherapy students 23.4%, 53 public health students 24.3%, and 53 basic medical sciences students 24.3%). As shown in Table 1.1, the majority of respondents were male (51.4%) followed by female (48.6%). Most of the respondents were aged with a mean score of 27.92 ± 3.195. Moreover, majority of respondents have experience less than 5years (75.2%) and (22.8%) has experience above than 5years. Value of perception and readiness in Shapiro-Wilk is .000 which shows that the data variable is not normally distributed. correlation of students towards interprofessional learning by applying Spearman’ rho test. Students’ perception has strong positive correlation with their readiness, p value (.000). Also students’ readiness has strong positive correlation with their perception towards interprofessional learning. This study was to explore the readiness and perception of students towards interprofessional education in different health care students. IPL is beneficial for students to know other professionals to work together for teamwork and collaboration and it also increase communication between different health care professional and patients.

## Introduction

Inter-professional learning (IPL) is an approach that teaches students diverse disciplines to communicate with each other about their professional knowledge in order to acquire a more complex one understanding of the current situation. According to WHO definition of IPE “it’s a process between two or more professionals learn about from, and with one another to permit effective association and enhance health outcome (1). Preparation of students for joint practice is one of the important role play by IPE. By providing meaningful opportunities for interaction with single professions to engage students with other disciplines, Interprofessional education allows students to reflect their own roles in multi-disciplinary team, know the roles of others and develop effective teamwork communication skills that are transferable to clinical practice(2). Students have to prepare before their professional life through inter-professional education and encouraging strategies by teachers regarding teamwork and collaboration, which will be helpful to become a more effective member of Health care team and achieve maximum health care outcome in patients living with chronic diseases. Students learn by sharing with other healthcare disciplines and better understand clinical problems. (3)It has been shown that medical errors can be reduced through improved interdisciplinary communication, which is one of the benefits of IPL, particularly when the learning groups have balanced input from each of the other professions. health professional students from discipline-focused programmes may have diverse attitudes and readiness towards participation in the IPL. The students’ preparedness to engage in IPL will be directly dependent on their attitudes and readiness. (4)The goal of IPE for students is to learn to function as part of an interprofessional team and to improve patient outcomes through interaction professional cooperation in your future practice. Greater coordination of health professionals through interdisciplinary collaboration has been shown to benefit patients prevent fragmentation of care and improve a holistic approach access to health. (5)One of the important aspects of interprofessional learning is concerned about changes in attitudes among professionals groups that may need research when considering shared learning. Thus Applying the principles of servant leadership can enhance the practice of professional development and strengthen the relationships between students in society, so that there is an increase respect the contributions and skills of different disciplines.(6). (7)Generally, for successful IPE programs positive attitude toward other professionals and team working is required. (8)The same understanding can howeveris not available to all team members. One of the report shows that medical students have the least confidence in their professional role and in their faith which requires more knowledge and skills than nursing or pharmacy students.(9). The aim of this study was to assess the perception and readiness towards inter-professional education among different health care disciplines at Khyber Medical University Peshawar. IPL helps students become familiar with key roles in their work, as well as the role of their team members from other professions. Recent research has shown that the value of previous IPL experience in the workplace lies in a better professional identity and attitude towards teamwork. Practice is important because the department seeks to provide a supportive learning experience cooperation of disciplines after graduation and into practice. Provide to the elderly an adult with experience in interdisciplinary teams improves the quality of their care, and providing students of different disciplines a real view of life in advanced age. This study addresses the degree of perception and readiness towards IPE among students of different healthcare disciplines. The study also helps the administrators to introduce and implement IPE in curriculum as well as its significant results also affect the quality of communication with patients.

## Methods and Material

### Study design

A correlational cross-sectional study design was conducted for the period of 4 months from September to December 2020, in different departments of Khyber medical university Peshawar. Convenience sampling was undertaken. The sample size was calculated using OpenEpi software by considering the total population of students during the study period; response distribution as 50% confidence interval was set at 95 and margin of error was at 5%. All the students were invited to participate in the study. The minimum sample size required tofulfill the study findings were a total of 218 students (61 Nursing students, 51 physiotherapy students, 53 public health students, and 53 basic medical sciences students completed the survey). Post graduate students from all discipline were included in the study. Undergraduates and students who were not interested in participating were excluded from the study.

### Setting, participants and ethical considerations

The study was conducted at Khyber medical university, Peshawar Pakistan, which provides healthcare related programmes. The sample was drawn from all postgraduate students enrolled in 2 years’ post-graduation degree program. Participation was voluntary. A written consent was obtained from the student prior to receiving the questionnaire. The study protocol was accepted by Khyber medical university ethical committee review board. Data obtained was stored safely for the specific period of time according to the requirements. Privacy and confidentiality will be maintained strictly. Risks and benefits ratio of the study was explained to the participants as per the requirement of the study.

### Study instruments

The Readiness for Interprofessional Learning Scale (RIPLS) developed by Parsell and Bligh (1999), which enabled the students to reflect on various aspects of IPL, was used to measure student readiness, or student beliefs, about IPL. A 5-point Likert Scale (Strongly agree = 5, agree = 4, neutral = 3, disagree = 2 and strongly disagree = 1) was used to analyze the students’ responses. The study tool has 19 self-reported items under four different domains. Domain 1 focused on the aspects of teamwork and collaboration (item 1–9). Domain 2 focused on positive and negative professional identity towards other professions (item 10–16). Domain 3 focused on the roles and responsibilities of professionals (item 17–19). The Interdisciplinary Education Perception Scale (IEPS)developed by Luecht, Madsen, Taugher, and Petterson (1990) was the second instrument used in the study to detect changes in learning over time among health professional students. It consisted of three domains (proficiency and dependence, Perceived Need for teamwork, and Perception of Actual Collaboration) with 18 items. The validated instrument used a 6-pointLikert-scale (Strongly disagree = 1, moderately disagree = 2, somewhat disagree = 3, somewhat agree = 4, moderately agree = 5 and strongly agree = 6). Additionally, the participants’ demographic details (age, gender, Programme of study and year of experience in field work) were also collected.

### Data Collection

Data collection were conducted in different departments of Khyber Medical University i.e. (INS, IPMR, IBMS, IPH&SS). Those respondents who have agreed and signed informed consent data were collected. Prior to signing the consent form students were informed about the confidentiality and given an explanation about the research aim. The procedures for completing the questionnaires were described after the questionnaire’s distribution. Students completed the questionnaires manually by ticking the statements which are match with their perceptions.

## Results

The RIPLS was completed by a total of 218 participants (response rate 100%, 61 Nursing students 28%, 51 physiotherapy students 23.4%, 53 public health students 24.3%, and 53 basic medical sciences students 24.3%). As shown in Table 1.1, the majority of respondents were male (51.4%) followed by female (48.6%). Most of the respondents were aged with a mean score of 27.92 ± 3.195. Moreover, majority of respondents have experience less than 5years (75.2%) and (22.8%) has experience above than 5years. (Fig 1.1-1.4)

Table 2 showing the significance value of perception and readiness in Shapiro-Wilk is .000 which shows that the data variable is not normally distributed. Just because the data is not normally distributed Pearson correlation test was applicable to use in this study. So we will use Spearman’ rho correlation test to check the correlation of student’s readiness and perception.

**Table 2.**
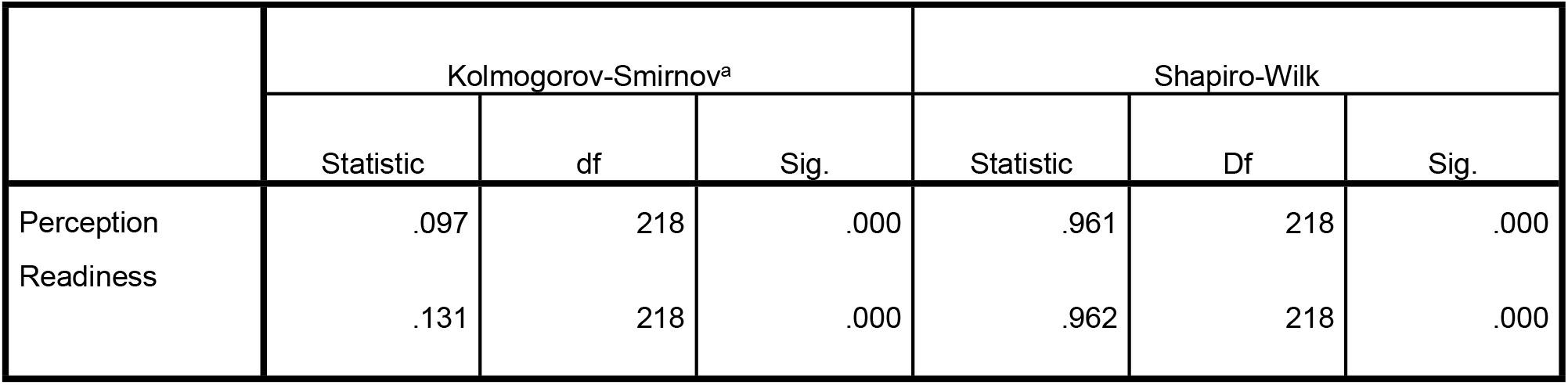
Tests of Normality.

Table 3 showing correlation of students towards interprofessional learning by applying Spearman’ rho test. Students’ perception has strong positive correlation with their readiness, p value (.000). Also students’ readiness has strong positive correlation with their perception towards interprofessional learning.

**Table 3.** Correlations b/w Students and IPL.

|  |  | Perception | Readiness |
| --- | --- | --- | --- |
| Perception | Pearson Correlation | 1 | .522** |
|  | Sig. (2-tailed) |  | .000 |
|  | N | 218 | 218 |
| Readiness | Pearson Correlation | .522** | 1 |
|  | Sig. (2-tailed) | .000 |  |
|  | N | 218 | 218 |

\*\* . Correlation is significant at the 0.01 level (2-tailed).

Table 4 showing the mean score for the readiness and perception. Mean score for the readiness of the students was 69.8 (SD= 8.41) and a minimum of 44 and maximum of 93 out of a possible 95. And the mean score for the perception students towards interprofessional education was 74.90 (SD=13.81) and a minimum score of 30 while a maximum score of 100 out of a possible 108.

**Table 4.** Mean score of Readiness and perception of all discipline.

|  | Perception | Readiness |
| --- | --- | --- |
| N | 218 | 218 |
| Mean | 74.9083 | 69.7982 |
| Std. Deviation | 13.81146 | 8.41743 |
| Minimum | 30.00 | 44.00 |
| Maximum | 100.00 | 93.00 |

Table 5 showing the results of each subscale for perception of the students towards inter-professional education for all discipline were: proficiency and dependence was 20.68 (SD 4.67) with a minimum of 7 and a maximum of 28. Score for subscale: perceived need for teamwork 16.38 (SD 3.39) with a minimum of 7 and a maximum of 23. Score for subscale: perception of actual collaboration was 20.40 (SD 4.33) with a minimum of 7 and a maximum of 28. Score for subscale: considering other values was 17.42 (SD 3.54) with a minimum of 7 and a maximum of 24. While the results of each subscale of readiness for all discipline were: Teamwork and collaboration; was 24.21 (SD 4.13) with a minimum of 10 and a maximum of 30. Score for subscale: Professional identity was 24.32 (SD 3.68) with a minimum of 15 and a maximum of 34. Score for subscale: Roles and responsibility was 8.80 (SD 2.25) with a minimum of 3 and a maximum of 15.

**Table 5.**
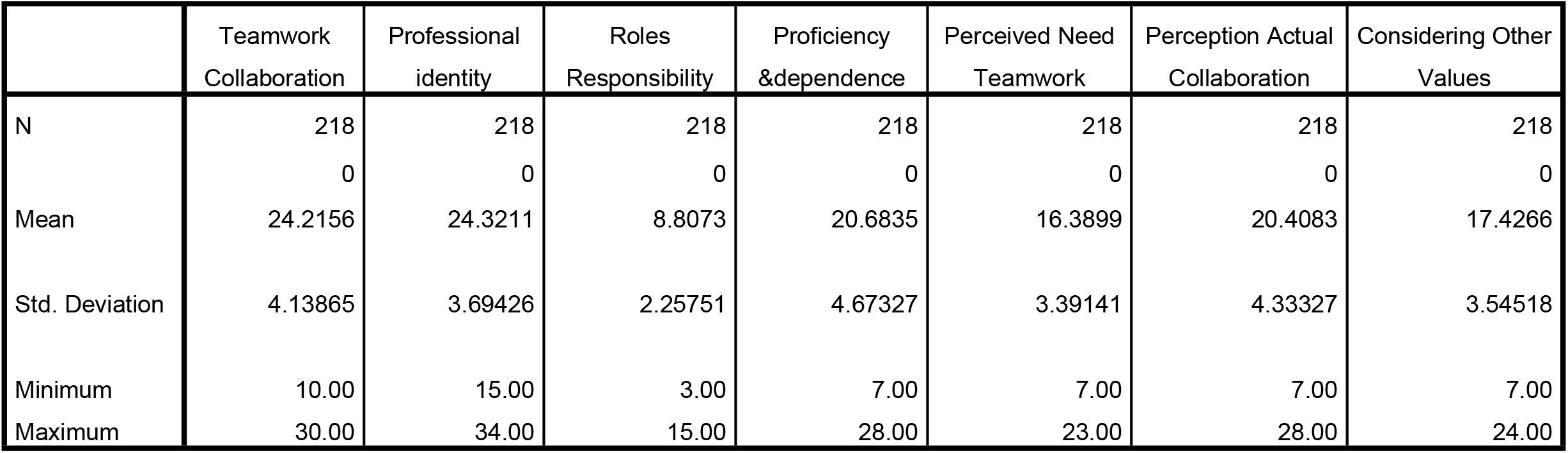
Subscale for IEPS & RIPLS among all disciplines.

|  | Teamwork<br>Collaboration | Professional<br>identity | Roles<br>Responsibility | Proficiency<br>&dependence | Perceived Need<br>Teamwork | Perception Actual<br>Collaboration | Considering Other<br>Values |
| --- | --- | --- | --- | --- | --- | --- | --- |
| N | 218 | 218 | 218 | 218 | 218 | 218 | 218 |
|  | 0 | 0 | 0 | 0 | 0 | 0 | 0 |
| Mean | 24.2156 | 24.3211 | 8.8073 | 20.6835 | 16.3899 | 20.4083 | 17.4266 |
| Std. Deviation | 4.13865 | 3.69426 | 2.25751 | 4.67327 | 3.39141 | 4.33327 | 3.54518 |
| Minimum | 10.00 | 15.00 | 3.00 | 7.00 | 7.00 | 7.00 | 7.00 |
| Maximum | 30.00 | 34.00 | 15.00 | 28.00 | 23.00 | 28.00 | 24.00 |

**Table 6.**
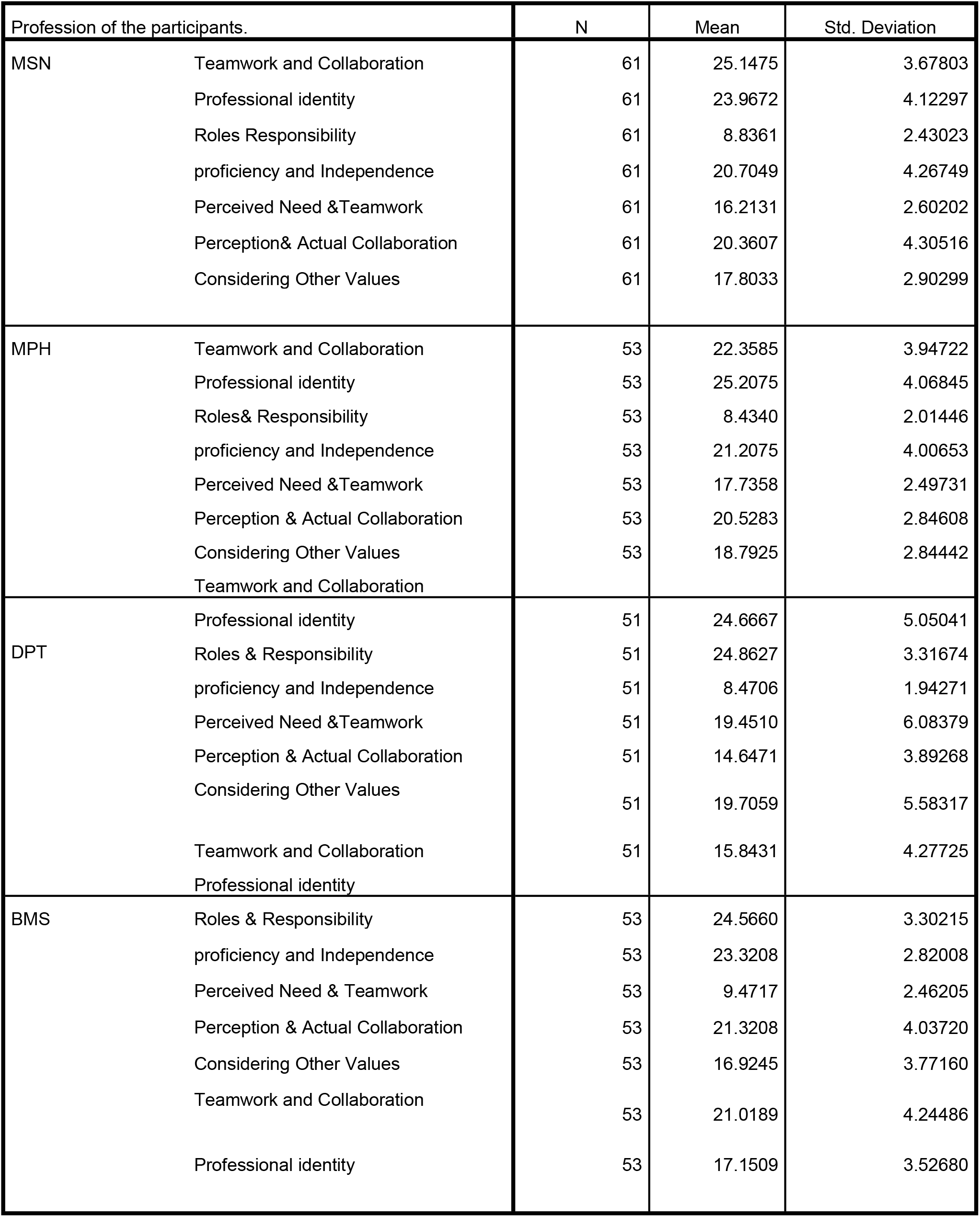

### Demographic data of the Participants

| Characteristics. | n=218 | Percentages. | Academic Program | n=218 | Percentages. |
| --- | --- | --- | --- | --- | --- |
| **Sex** |  |  | Nursing | 61 | 28% |
| Male | 112 | 51.4% | Physiotherapy | 51 | 23.4% |
| Female | 106 | 48.6% | Public Health | 53 | 24.3% |
| **Age** |  |  | Basic Medical Sciences | 53 | 24.3% |
| <30 | 163 | 74.8% |  |  |  |
| >30 | 55 | 25.2% |  |  |  |
| **Experience** |  |  |  |  |  |
| <5 Years | 164 | 75.2% |  |  |  |
| >5 Years | 54 | 24.8% |  |  |  |

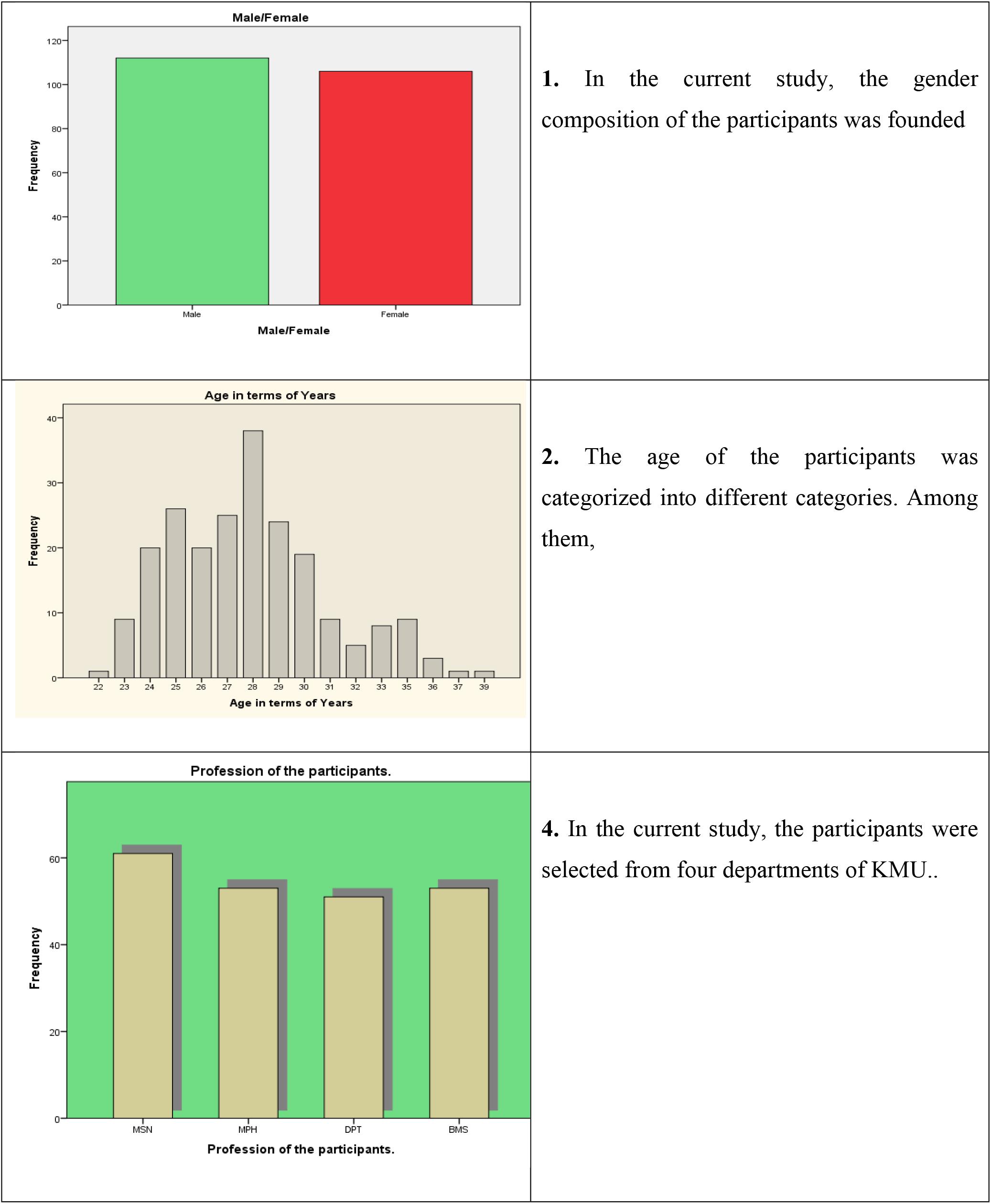
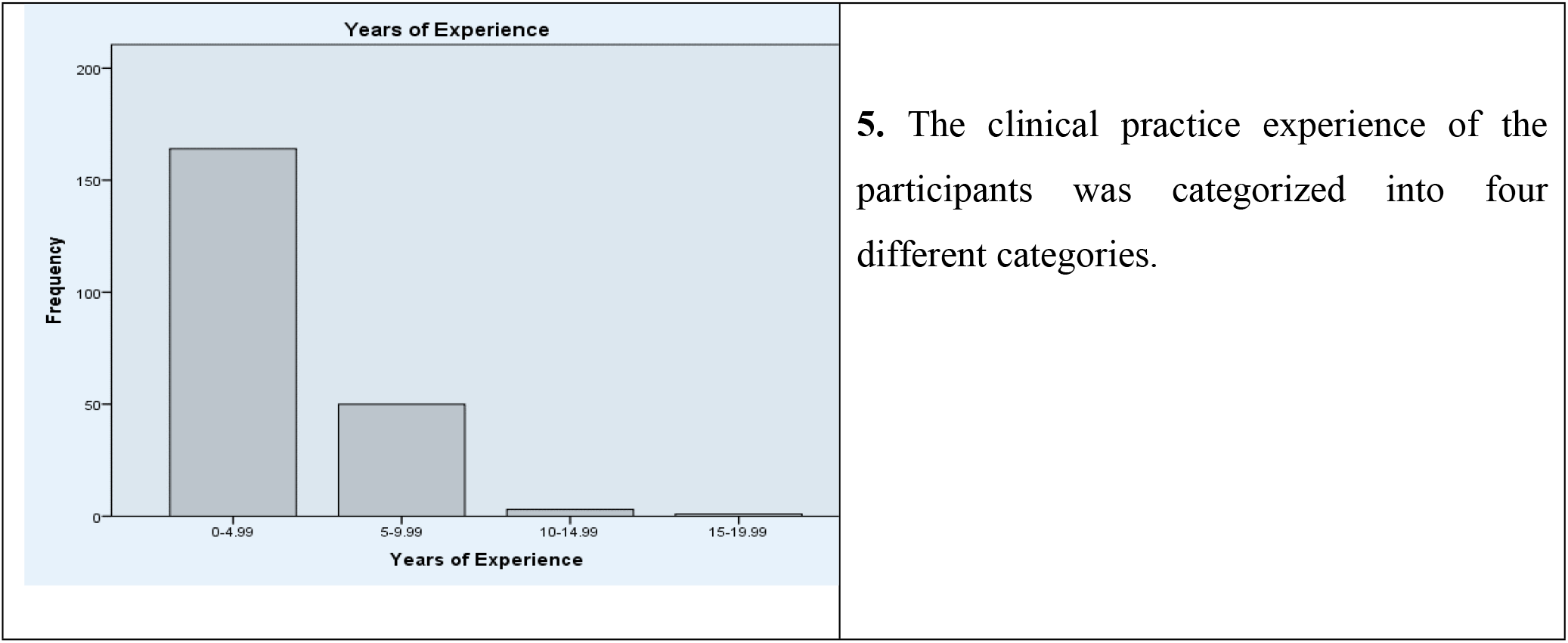

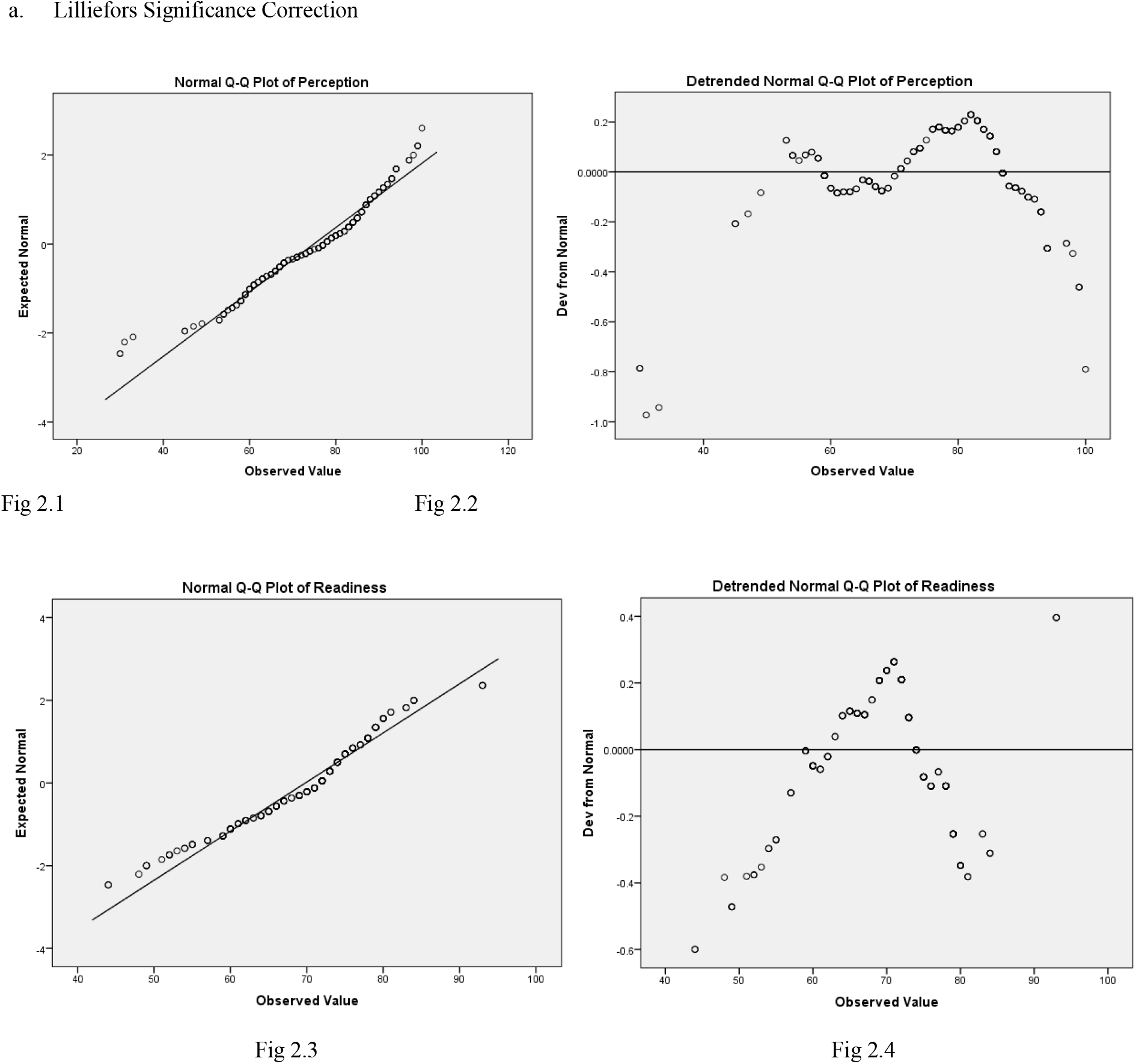

## Discussion

The aim of this study was to explore the readiness and perception of students towards interprofessional education in different health care students. IPL is beneficial for students to know other professionals to work together for teamwork and collaboration and it also increase communication between different health care professional and patients. The result is that students can learn professional skills from each other and enhanced understanding between them as well as patients’ problem. While the advantage of interdisciplinary education among different healthcare students is to understand their professional role and allow them to be able to work closely with individuals in other profession. (10). The participants of this study was postgraduation students who were enrolled in a 2 years’ degree program in Khyber Medical University Peshawar from different disciplines that include Master in Nursing 28%, Master in physiotherapy 23.4%, Master in public health 24.3% and Master in basic medical sciences 24.3% i.e. (Anatomy, Microbiology, Biochemistry, Pharmacology and histopathology) 24.3% in which 51.4% male and 48.6% female. Study distribution was check by applying Shapiro-Wilk W test to check the normality of the data. As the result was 0.000 and it was less than 0.05 so it was statistically significant that the sample was not normally distributed and we reject the null hypothesis that the sample was drawn from a normal distribution. Spearman’s rho was correlation test was used to check the correlation of students’ perception and readiness. Participants shows strong positive correlation in their readiness and perceptions towards interprofessional learning, (p=.000). the results are in agreements with some previous studies, which shows most healthcare students have positive perception and readiness towards IPL in undergrads level. (1). Our results show that on the subscale teamwork and collaboration majority of students were agreed with importance of collaboration and teamwork domain with other health care professionals. This result matches the previous finding study that observed students had positive attitude towards teamwork and collaboration.(9). The high values of our results in professional identity subscale for the first three questions that show negative professional identity in this domain highly suggests by all disciplines students that for undergrads it is not necessary to learn together with other healthcare professional and clinical problem solving skills can be learned from my own department students’ as well as also had a high score of students’ agreement from each health profession group agreed or strongly agreed with the four items in this section that show positive professional identity. Respondents from all disciplines agreed that shared learning with other health care professionals would help them to communicate better with patients and other professionals. The item in subscale roles and responsibility are concerned with the idea that professional clinical practice improves health professionals’ responsibility. Results to the following question in this domain “the function of the nurses and therapists is mainly to provide support for doctors” response of MSN students was 62.29% strongly disagree followed by Basic medical sciences students 56.60%, Public health students 52.83% and physiotherapy students 47.05%. Our study findings were collected through a valid and reliable questionnaire. It is an appropriate instrument that measure students’ readiness towards IPE.The RIPL scale was validated for middle eastern populations. The authors reported the instrument as having high content validity and an internal consistency reliability of 0.9. (11). The data collected from 218 students represented a response rate of 100% and it was close to the recommended sample size. Our study findings may be useful for students who are being considered as future members of rehabilitation teams, faculty members of universities, and rehabilitation teams in health-care centers.

Some of the challenges encountered in our study affected the result. The short period of time restricted the involvement of our health profession education students in our study; therefore, we are unable to include all the disciplines related to health care profession team. The subject was selected from one university of Peshawar therefore the results cannot be generalized to students of other universities or contexts. This was a cross sectional study design that is not able to exactly determine the factors that affect the differences among the results. Ideally, IPE would promote specific learner skills, including teamwork, leadership, consensus building and the ability to recognize achieve common goals of patient care. Course administration is an interprofessional teamwork in the health program a great challenge for the clinical teaching community. IPE offers the opportunity to address the multidisciplinary concept through hospitals.

## Conclusion

Present study findings showed strong positive correlation between perception and readiness for interprofessional learning. Positive attitude of students towards interprofessional training of qualified health professionals to note, share and learn from each other new skills and practices experience and knowledge. The most important findings of this study was subscale of roles and responsibility domain question “the function of the nurses and therapists is mainly to provide support for doctors” response of MSN students strongly disagree followed by Basic medical sciences students, Public health students, and physiotherapy students. Interprofessional education is very important and valuable to develop a willingness to work with others who health at work is a very important asset especially realizing what science has done today cooperative health cooperative discussion.

